# Decreasing hearing ability does not lead to improved visual speech extraction as revealed in a neural speech tracking paradigm

**DOI:** 10.1101/2024.03.13.584400

**Authors:** Chandra Leon Haider, Anne Hauswald, Nathan Weisz

**Author notes:** corresponding author Email address: Chandra Leon Haider Anne Hauswald, Nathan Weisz. The authors have declared no competing interests.

## Abstract

The use of visual speech is thought to be especially important in situations where acoustics are unclear and in individuals with hearing impairment. To investigate this in a neural speech tracking paradigm, we measured MEG in sixty-seven mid- to old-age individuals during audiovisual (AV), audio-only (A), and visual-only (V) speech in the context of face masks. First, we could extend previous findings by showing that not only in young normal-hearing individuals but also in aging individuals with decreasing hearing ability the brain is superior in neurally tracking the acoustic spectrogram in AV compared to A presentations, especially in multi-speaker situations. The addition of visual lip movements further increases this benefit. Second, we could show that neural speech tracking in individuals with lower levels of hearing ability is affected more by face masks. However, in this population, the effect seems to be a composite of blocked visual speech and distorted acoustics. Third, we could confirm previous findings, that the neural benefit of visual speech varies strongly across individuals. We show that this general individual ability predicts how much people engage in visual speech tracking in difficult AV listening situations. Interestingly, this was not correlated with hearing thresholds and therefore does seem to be a widely used compensatory strategy in the hearing impaired.

## Introduction

Successful speech processing and production are the most important components of everyday communication. In terms of perception, it is therefore not surprising that humans do not rely on a single sensory modality such as audition. As speech production inherently also contains a visually observable component, the speech recipient can use it to help with speech perception. Early studies have already explored how visual information helps with comprehension (Sumby & Pollack, 1954), but also how incongruent visual information can lead to a wrong perception of the audio input has been demonstrated by the McGurk effect (Mcgurk & Macdonald, 1976). Past theories proposed mechanisms for how visual speech is used to enhance speech processing (Peelle & Sommers, 2015). On the one hand, in an early processing stage, visual speech provides cues on when to focus attention on the auditory input. On the other hand, in a late processing stage, visual and auditory inputs are combined for phoneme discrimination. Similarly, to the McGurk effect, some phonemes are acoustically ambiguous and can more clearly be categorized with the help of visual speech. Alongside the growing understanding through behavioral studies, also underlying neural mechanisms have been explored. In recent years, the possibility arose to analyze the brain’s response to continuous speech inputs, which is a much more naturalistic stimulus compared to short sentences that have been used in classic ERF/ERP analyses (e.g. Besle et al., 2004, 2009). Among various options (e.g. coherence and cross-correlation), so-called temporal response functions (TRFs) have recently gained popularity. (Brodbeck et al., 2022; Crosse, Di Liberto, Bednar, et al., 2016). Through this, it is possible to obtain a measure that can be viewed as the strength of the representation of a certain speech characteristic in the brain. With the help of these and similar techniques several studies could show general processing of visual-only speech (V), but also how audiovisual (AV) compared to audio-only (A) speech improves the representation of the speech input in the brain (Crosse, Di Liberto, & Lalor, 2016; Crosse et al., 2015; Hauswald et al., 2018; O’Sullivan et al., 2021; Park et al., 2016; Suess et al., 2022).

In three past magnetoencephalography (MEG) speech tracking studies, we highlighted the importance of visual speech (Haider et al., 2022, 2023; Reisinger et al., 2023). In Haider et al. (2022), we showed how the reconstruction of different speech features is affected when masking the mouth area with a face mask in clear and multi-speaker speech. We followed it up in Haider et al. (2023) showing on the one hand, that the adverse effects of the face mask are indeed visual and not the product of distorted acoustics, on the other hand, we could highlight the benefits of visual speech, especially their importance in multi-speaker situations as well as topographically localizing them. We demonstrated that by, on the one hand, calculating the visual benefit as the difference between AV and A conditions, and on the other hand calculating the added value through lip movements. Finally, by using the same data as Haider et al. (2022), Reisinger et al., (2023) linked neural and behavioral findings. They showed that individuals who profit more from lip movements on a neural level show worsened behavioral performance when the speaker is wearing a face mask, thereby further strengthening the idea of inter-individual differences in how individuals rely on visual speech information (Aller et al., 2022).

However, all of our studies are limited in that they only investigated young and normal-hearing individuals. These participants are expected to be less affected by degrading the speech input acoustically as compared to individuals suffering from hearing impairments. In fact, individuals with hearing impairments were reported to be affected most detrimentally by the widespread use of face masks during the Covid-19 pandemic (Brown et al., 2021; Chodosh et al., 2020; Homans & Vroegop, 2021). Additionally, a study by Puschmann et al. (2019) showed stronger visual benefits in speech in individuals reporting high listening effort compared to individuals reporting low effort. However, they only focussed on the reconstruction of the speech envelope. With the same paradigm as in Haider et al. (2023), we now want to investigate how and if individuals with a wide range of hearing abilities differ in how they profit from visual speech. To do that, we presented individuals with A and AV speech. Furthermore, in half of the conditions, the speakers wore a face mask, while a second distractor speaker was added in half of the conditions. With that, we can differentiate how individuals with different levels of hearing ability profit from visual speech and if this changes in multi-speaker compared to single-speaker listening situations. We also explore which brain regions and associated mechanisms are employed for extracting relevant visual speech components. Finally, by using V speech we find hints for a measure that might be a general predictor of how much they employ visual speech in their speech processing.

## Methods

### Participants

67 German native speakers (36 female) aged between 41 and 72 years (M = 59.97, SD = 5.55) took part in our study. The exclusion criteria were non-removable magnetic objects, as well as a history of psychiatric or neurological conditions. To assess the individual hearing ability, we computed the average hearing threshold of both ears and across all frequencies using standard clinical audiometry. Both, age and hearing ability are more thoroughly presented in Supplementary Material S1. We recruited participants as part of the *Paracelsus 10,000* (Frey et al., 2023) assessment or recruited via our subject database. All participants signed an informed consent form. The experimental protocol was approved by the ethics committee of the University of Salzburg and was carried out in accordance with the Declaration of Helsinki.

### Stimuli

For this study, we used the same stimulus material as in Haider et al. (2023). We used excerpts from four different stories for our recording read out in German. ‘Die Schokoladenvilla - Zeit des Schicksals. Die Vorgeschichte zu Band 3’ (“The Chocolate Mansion, The Legacy” – a prequel of Volume 3”) by Maria Nikolai and ‘Die Federn des Windes’ (“The feathers of the wind”) by Manuel Timm were read out by a female speaker. ‘Das Gestüt am See. Charlottes großer Traum’ (“The stud farm by the lake. Charlotte’s great dream”) by Paula Mattis and ‘Gegen den Willen der Väter’ (“Against the will of their fathers”) by Klaus Tiberius Schmidt were read out by a male speaker.

Stimuli were recorded using a Sony FS100 camera with a sampling rate of 25 Hz for video and a Rode NTG 2 microphone with a sampling rate of 48 kHz for audio. We aimed at a duration for each story of approximately ten minutes, which were cut into ten videos of around one minute each (range: 56–76 s, M = 63 s, SD = 5.0 s). All stories were recorded twice, once without the speaker wearing a surgical face mask and once with the speaker wearing a surgical face mask (Type IIR, three-layer single-use medical face mask, see Fig. 1A). After cutting, all videos were approximately one minute in length (80 AV recordings in total). Thirty of those were presented to each participant (15 with a female speaker, 15 with a male speaker) in order to rule out the sex-specific effects of the stimulus material. The audio track was extracted and stored separately. The audio files were then normalised using the Python function ‘ffmpeg-normalise’. Pre-recorded audiobooks read out by different speakers (one female, one male) were used for the distractor speaker and normalised using the same method. The syllable rate was analysed using a Praat script (Boersma and Weenink, 2001; de Jong and Wempe, 2009). The target speakers’ syllable rates in the 30 trials varied between 3.65 Hz and 4.57 Hz (M = 4.04 Hz, SD = .25). Target and distractor stimuli were all played to the participant at the same volume, which was individually set to a comfortable level at the start of the experiment by using an example audiobook with the target female speaker.

**Fig. 1.**
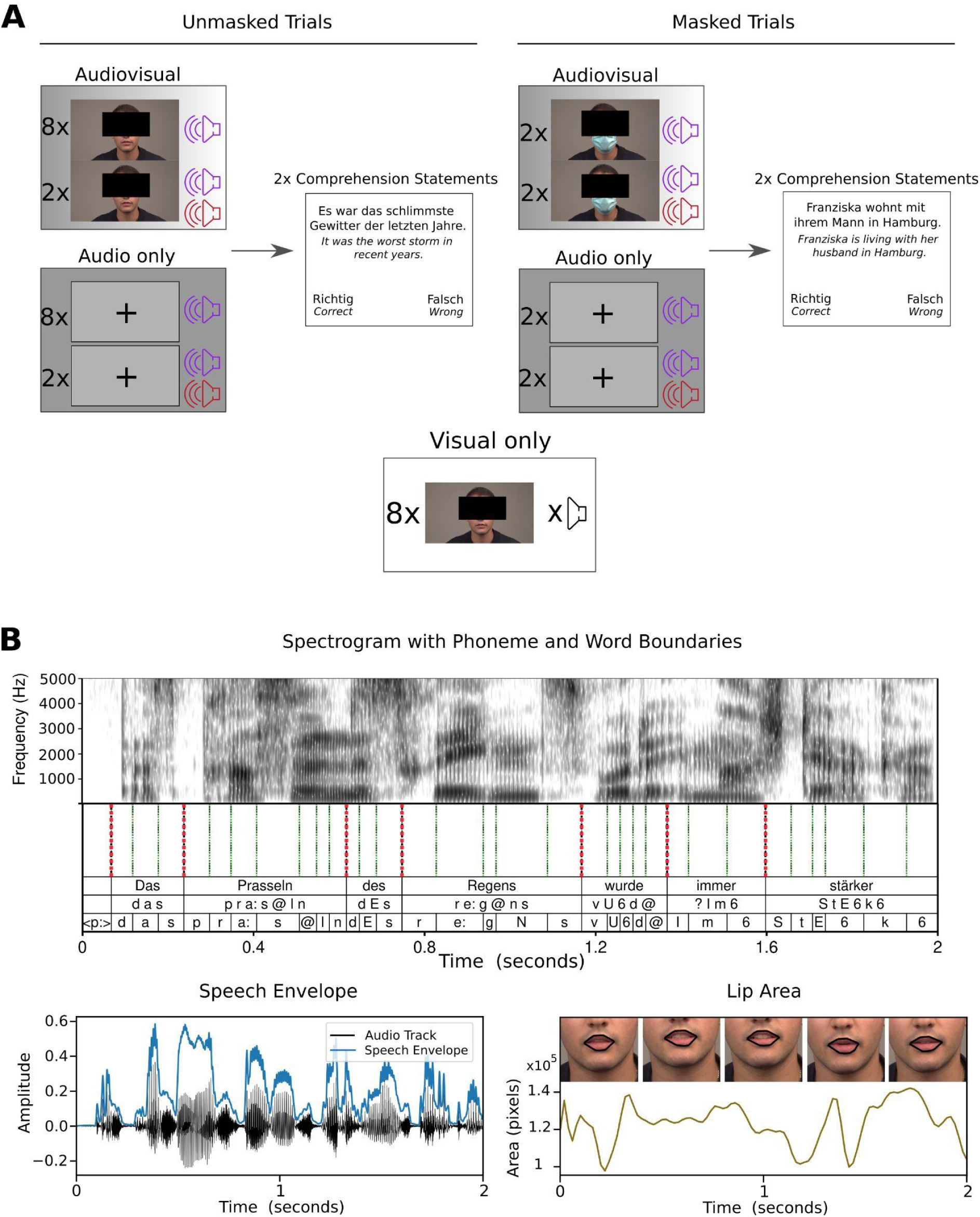
Experimental procedure and speech features. **A** The condition design for the experiment with the male speaker as an example. Every participant was presented with 36 trials of ∼1 min each. We conducted AV trials (14 trials in total), A trials (14 trials in total), and V trials (8 trials in total). For A and AV trials, we used 10 unmasked trials each. In eight of these trials, only a single target speaker was presented. In two of those trials, we added a second same-sex (distractor) speaker (denoted by the second sound icon). We used four masked trials for A and AV speech, respectively. In two of those trials, again only a single target speaker was presented, and in two trials, we added a distractor speaker. The design was unbalanced to obtain sufficient training data to compute the temporal response function models (see TRF Model Fitting section). After each A and AV trial, we prompted two “true or false” statements to measure comprehension and to keep participants focused. Participants answered via a button press (left or right button). **B** The investigated speech features. The spectrogram is shown alongside the investigated speech units, phonemes, and words underneath (top row: orthographic word; middle row: phonetic word; bottom row: phoneme). The speech envelope can be seen on the bottom left of the figure. On the bottom right of the figure, the extracted lip area can be seen with the presentation of the mouth outline corresponding to the 1st, 13th, 25th, 38th, and 50th frames (i.e., 0, 0.5, 1, 1.5, and 2 sec). All depictions are based on the same 2-sec-long speech interval. The spectrogram and speech envelope are depicted before downsampling for illustrative purposes.

### Experimental Procedure

For this study, we employed the same experimental paradigm as in Haider et al. (2023). Before the start of the experiment, we performed standard clinical audiometry between .125 and 8 kHz using an AS608 Basic (Interacoustics, Middelfart, Denmark) to assess participants’ hearing ability. We started the MEG measurement with five minutes of resting-state activity (not included in this manuscript). We then adjusted the stimulation volume by presenting an exemplary audiobook and adjusted the volume until participants stated that it was clearly audible and comfortable. One block consisted of six trials of ∼ 1 min length. The condition design is depicted in Figure 1. As an extension to Haider et al. (2022), we added audio-only modality (A) and visual-only modality (V) conditions in addition to the audiovisual modality conditions (AV). For the A trials, we used the same stimuli as the AV trials but did not show the corresponding video and instead presented a fixation cross. This results in 14 AV, 14 A, and 8 V trials. For A and AV conditions respectively, we still wanted to keep the 2x2 design from Haider et al. (2022). Therefore, 8 trials per modality consisted of clear speech (i.e. no mask and no distractor), while in the trials speakers wore a face mask, a second same-sex speaker was added or both (i.e. two trials per one of these conditions). This design was chosen to have sufficient data to train the TRF models on the one hand and to have a reasonable experiment duration on the other hand. All conditions had an equal amount of female and male speaker trials. In conditions with a second speaker (distractor), the second speaker only started five seconds after the first (target) speaker to give participants time to focus on the to-be-attended speaker. Within the first 4 blocks, the story presentation followed a consistent storyline across trials. The fifth and sixth block was a mixed speaker block (both female and male speakers) containing continuations from the former stories. Three trials each were used from ‘Gegen den Willen der Väter’ (male speaker), ‘Die Federn des Windes’ (female speaker) for the fifth block, and ‘Die Schokoladenvilla - Zeit des Schicksals. Die Vorgeschichte zu Band 3’ (female speaker) and ‘Das Gestüt am See. Charlottes großer Traum’ (male speaker) were used in the sixth block. Each story segment was only presented once per participant. Except for the V-only trials, two unstandardised ‘true or false’ statements regarding semantic content were asked at the end of trials to assess comprehension performance and keep participants focused (Fig. 1A). Additionally, participants rated subjective difficulty twice per condition on a five-point Likert scale for A and AV trials. The participants’ answers were given via button presses. The condition design was shuffled across trials to rule out any influence of the specific stimuli material on the results. As an exception to this, we always assigned a trial without a mask and without a distractor as the starting trial of each block to let participants adapt to a (possible) new target speaker. Videos were back-projected on a translucent screen with a screen diagonal of 74 cm via a Propixx DLP projector (Vpixx technologies, Canada) ∼ 110 cm in front of the participants. It was projected with a refresh rate of 120 Hz and a resolution of 1920 × 1080 pixels. Including preparation, the experiment took about 2 h per participant. The experiment was coded and conducted with the Psychtoolbox-3 (Brainard, 1997; Kleiner et al., 2007; Pelli, 1997) with an additional class-based library (‘Objective Psychophysics Toolbox’, o_ptb) on top of it (Hartmann and Weisz, 2020).

### Speech feature extraction

All the speech features investigated are depicted in Fig. 1B. The same features and extraction procedures were used as in Haider et al. (2023). The speech envelope was extracted using the Chimera toolbox. By using the default options, the speech signal was filtered forward and in reverse with a 4th order Butterworth bandpass filter at nine different frequency bands equidistantly spaced on the cochlear map between 100 and 10000 Hz (Smith et al., 2002). Then, a Hilbert transformation was performed to extract the envelopes from the resulting signals. These nine bands were then summed up to one general speech envelope and normalized. Finally, a 80 Hz low pass filter was applied on the envelope.

The spectrogram was computed similarly. We only adjusted the lower end of the frequency range to 50 Hz, as this then also includes the speech pitch / fundamental frequency (especially for the male speaker). In the end, the resulting nine bands were not summed up but used together as the speech spectrogram.

Phonemes and word onset values were generated using forced alignment with MAUS web services (Kisler et al., 2017; Schiel, 1999) in order to obtain a measure for speech segmentation. We generated two time-series at 50 Hz with binary values indicating an onset of a phoneme or word, respectively. Furthermore, we extracted every phoneme separately. In this case, we ended up with 63 time series, one for each phoneme, containing only zeros and ones.

The lip area was extracted from the video recordings using a Matlab function by Park et al. (2016). Through this, we generated a time series of the lip area in pixels with a sampling rate of 25 Hz (i.e. the frame rate of the video recording).

In the end, all features were sampled to 50 Hz to match the sampling rate of the corresponding brain signal, as most speech-relevant signals present themselves below 25 Hz (Crosse et al., 2021).

### Data acquisition

We recorded brain data with a sampling rate of 1 kHz at 306 channels (204 first-order planar gradiometers and 102 magnetometers) with a Triux MEG system (MEGIN, Helsinki, Finland). The acquisition was performed in a magnetically shielded room (AK3B, Vacuumschmelze, Hanau, Germany). Online bandpass filtering was performed from 0.1 Hz to 330 Hz. Before the acquisition, cardinal head points (nasion and pre-auricular points) were digitized with a Polhemus FASTRAK Digitizer (Polhemus, Colchester, Vermont, USA) along with around 300 points on the scalp to assess individual head shapes. Using a signal space separation algorithm provided by the MEG manufacturer (Maxfilter, version 2.2.15), we filtered noise resulting from sources outside the head and realigned the data to a standard head position, which was measured at the beginning of each block.

### MEG preprocessing

The raw MEG data was analyzed using Matlab R2020b (The MathWorks, Natick, Massachusetts, USA) and the FieldTrip toolbox (Oostenveld et al., 2011). As part of our standard pipeline, we first computed 50 independent components to remove eye and heart artifacts. We removed on average 2.88 components per participant (SD = 1.54). We further filtered the data using a sixth-order zero-phase Butterworth bandpass filter between 0.1 and 25 Hz. Finally, we downsampled our data to 50 Hz for more efficient computation while still preserving sufficient information from our data (Crosse et al., 2021).

### TRF model fitting

Fitting the temporal response functions (TRF) was identical to Haider et al. (2023) with the addition of the visual-only (V) conditions. In order to prepare the data for fitting the TRF models, we first z-scored the MEG data of all 306 channels and rescaled the speech envelope and lip area to values between 0 and 1. This rescaling was preferred over z-scoring, as this does not introduce negative values into only positive sign features. We used the mTRF Toolbox (Crosse, Di Liberto, Bednar, et al., 2016) to reconstruct stimulus features (in the case of backward modeling) or to predict brain data (in the case of forward models). For a more detailed discussion about this method please refer to Crosse et al. (2016) and Crosse et al. (2021). We trained separate models on audiovisual (AV) clear speech (i.e. AV speech without a face mask and without a distractor), on audio-only (A) clear speech, and on visual-only speech (V) respectively. We then used the A and AV to reconstruct the stimulus feature or predict brain activity for their respective modality (AV model for AV conditions and A model for A conditions) but across other conditions (i.e. for trials with masks and with distractors). As the second distractor speaker only starts after 5 seconds into the trial, we removed the first 5 seconds of each distractor trial from the analysis. V models were trained on V conditions and tested on V conditions to obtain a general score of visual speech tracking. We also tested the V model on AV conditions to compute a tracking value specifically tuned to visual speech but in a more natural AV situation.

For both forward and backward modeling, we used time lags from −50 to 500 ms to train our models. The regularization parameter (lambda) between 10^-6^ … 10^6^ was determined using a seven-fold leave-one-out cross-validation (Willmore & Smyth, 2003).

In order to completely rule out the possibility that a single outlier trial influences the model training, we again made use of a leave-one-out procedure. We used seven clear speech trials for model training and the eighth clear speech trial for model testing. We then looped across the trials so that every trial was once used for testing and otherwise used for training. In the end, we averaged the Fisher-z-transformed values across each iteration together. Overall, we used ∼ 7 minutes for training our A and AV models and ∼ 1 of testing for the clear speech condition in each iteration. For all other three conditions (i.e. No Mask + Distractor; Mask + No Distractor; Mask + Distractor), we used all trials for testing which was ∼ 2 minutes each.

#### Backward modeling

With this approach, we are able to map the brain response back to the stimulus feature (e.g. the speech envelope) to acquire an omnibus measure of how well a certain feature is encoded in the brain. This model has the benefit that the whole brain activity is taken into account when reconstructing a feature, which holds information about a general representation of a certain stimulus feature in the brain. By this approach, a single correlation value (Pearson’s r) is obtained as the measure of how well a feature can be reconstructed, which makes it easily interpretable. This however comes with the downside that the reconstructed features are all modeled independently of each other. Therefore, the amount of shared information between the features is not taken into account. It is therefore more commonly used on one-dimensional features (e.g. speech envelope) than on multi-dimensional features (e.g. spectrogram) as in the latter case each dimension is reconstructed independently, which makes claims on how well a certain feature is tracked as a whole difficult. Furthermore, by using this approach spatial information on which brain regions are involved in the processing of certain stimulus characteristics is lost. To solve both of these issues, one can use forward modeling as a complementary analysis.

#### Forward modeling

By using this approach, we are mapping the stimulus (-feature) ‘forward’ to the individual MEG channels. We therefore acquire a measure of how well a certain stimulus feature is encoded in the individual channels. In contrast to backward models, this model cannot give a measure of how well a feature is generally encoded in the whole brain, as it does not account for intercorrelations between the channels. However, as we get a prediction accuracy measure for each individual channel, it is possible to acquire a spatial distribution in the brain and test hypotheses about the contribution of different brain regions. Additionally, we can use multivariate inputs to predict brain data (e.g. spectrogram) which allows us to evaluate more complex (multidimensional) features. By adding a feature to a baseline model and subsequently comparing the new model to the baseline, one is able to obtain the contribution of the added feature above the simpler model. This is especially important in speech research, as speech features are highly correlated.

We did not analyze the resulting prediction accuracies on channel-level but beamed them to source-space.

### Source projection of MEG channels

After predicting our channel-level data via forward modeling, we used beamformers to project both the original and predicted channel data into the underlying source space. In order to do that we used the headmodels of the participant obtained through the digitization of the participant’s head. Then we used a 3D grid and morphed the participants’ heads onto it by utilizing a standard structural brain sourced from the Montreal Neurological Institute (MNI) in Canada. The latter was then adjusted to accurately align with the individual’s fiducials and head shape, following the methodology outlined by Mattout et al. in 2007. Subsequently, a 3D grid with 1 cm resolution and 2982 voxels, based on an MNI template brain, was morphed to conform to the specific brain volume of each participant. This alignment facilitates group-level averaging and statistical analysis, as all grid points within the transformed grid correspond to the same brain region across subjects.

These aligned brain volumes were also employed in generating single-shell head models and lead fields, following the approach outlined by Nolte in 2003. By utilizing the lead fields and a common covariance matrix (which aggregates data from all blocks), a shared LCMV beamformer spatial filter was computed, as described by Van Veen et al. in 1997.

We used a brain atlas (Tzourio-Mazoyer et al., 2002) to remove the cerebellum and unlabeled regions (primarily medial white matter, brain stem, and ventricles) from the grid. By doing that out of the 2982 voxels 1266 remained.

### Statistical analysis

We examined the forward models using one-tailed dependent sample cluster-based permutation tests, employing 10,000 permutations according to the methodology outlined by Maris and Oostenveld (2007). The *maxsum* method, integrated into Fieldtrip, was used for computing the t-statistic. Additionally, we assessed the effect size of each cluster by averaging Cohen’s d values across all significant channels within the cluster. The selection of a one-tailed test was based on our investigation of differences between simple and additive models, with the expectation that the performance of the latter should be at least equivalent to that of the former. We performed this analysis by using the remaining 1266 voxels as described in the *Source projection of MEG channels* section.

For analyzing the backward models as well as the behavioral data we used a Bayesian multilevel model approach with the *brms* package (Bürkner, 2017). We averaged the data for each condition and each dependent variable of interest. We included a random intercept for each subject as well as random slopes for each of our experimental manipulations (i.e. *Mask* (0/1), *Multispeaker* (0/1), and *Modality* (A/AV)).

𝑦 ∼ 1 + 𝑀𝑎𝑠𝑘 * 𝑀𝑢𝑙𝑡𝑖𝑠𝑝𝑒𝑎𝑘𝑒𝑟 * 𝑀𝑜𝑑𝑎𝑙𝑖𝑡𝑦 * 𝐻𝑒𝑎𝑟𝑖𝑛𝑔𝐴𝑏𝑖𝑙𝑖𝑡𝑦 +

(1 + 𝑀𝑎𝑠𝑘 + 𝑀𝑢𝑙𝑡𝑖𝑠𝑝𝑒𝑎𝑘𝑒𝑟 + 𝑀𝑜𝑑𝑎𝑙𝑖𝑡𝑦 | 𝑆𝑢𝑏𝑗𝑒𝑐𝑡)

with y being the dependent variable of interest (e.g. speech envelope or comprehension scores). We modeled the data with a Student-t distribution. Pareto k values revealed a good fit of the distribution.

## Results

First, we are presenting the added value of lip movements and lexical units to a baseline acoustic model in source space similar to Haider et al. (2023). Then, we follow this up with the results regarding the effects of hearing ability on neural speech tracking and behavioral performance. Finally, we conclude this section with results regarding interindividual differences in visual speech tracking and its impact in multi-speaker listening situations.

### Source reconstruction reveals visual speech benefits in occipital regions

With this analysis, we replicated the approach from (Haider et al., 2023) in source space in a new and independent sample. By using the mTRF Toolbox in the forward direction (Crosse, Di Liberto, Bednar, et al., 2016), we calculated one correlation coefficient per condition, per channel, and per subject for each investigated feature. As speech features are proven to be strongly intercorrelated, the use of additive models is highly beneficial to investigate differences and potential gains by adding additional features to the models. Thereby, it is possible to differentiate the unique variance explained by the added feature. For this analysis, we computed a (base) acoustic model (spectrogram), a spectrogram + lip model (spectrogram + lip area), and a spectrogram + lexical model (spectrogram + phonemes + word onsets). The resulting correlation coefficients from the models were Fisher-z transformed and averaged within conditions. To investigate the effects statistically, we used a one-tailed dependent-sample cluster-based permutations test, as described in the Methods section. The results are presented visually in Figure 2. By *added value*, we refer to increases in tracking through adding features beyond the spectrogram to predict the brain data (i.e. Spectrogram + Lip). By *visual benefit*, we refer to increases in tracking in the AV condition compared to the A condition.

**Figure 2.**
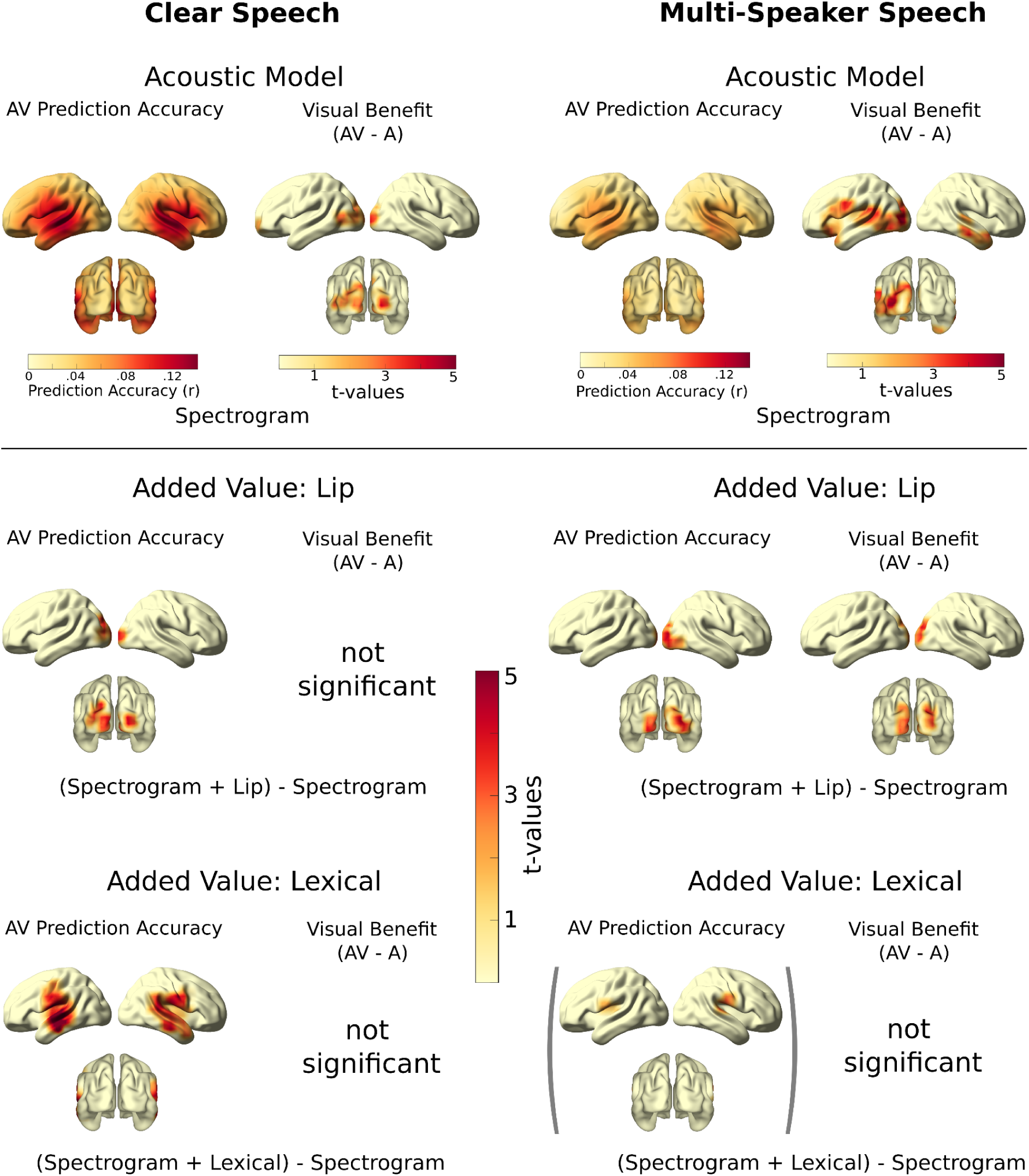
Illustration depicting the outcomes of forward modeling and its added value. The top row showcases the prediction accuracy and visual enhancement across the brain surface using the baseline acoustic model based on the spectrogram. Data on the left derive from participants exposed to clear speech, while data on the right pertain to those exposed to multi-speaker (distractor) speech. A statistical examination of the visual enhancement (i.e., AV–A) for the acoustic model was conducted. The subsequent rows depict the added value through lip movements (lip area) and lexical features (phonemes and word onsets), respectively. Specifically, they denote the difference between the spectrogram model and the combined (complex) model (spectrogram + lip or spectrogram + lexical) concerning ‘raw’ prediction accuracies and visual benefit. For the contrasts, significant clusters (p < .05, one-tailed) are shown. One trend (p < .1, one-tailed) for the added value of the lexical model in distractor speech is shown in parentheses.

#### Spectrogram

For the acoustic baseline model (as depicted in the top row of Figure 2), we investigated the differences between A and AV conditions in clear and multi-speaker speech. In clear speech, the results show two significant clusters. As expected, one large cluster was located in occipital areas (t_sum_ = 220.52, p = .002, d = .68) showing increased tracking in AV compared to A speech. A second positive cluster was located in left frontal regions (t_sum_ = 84.53, p = .031 d = .40). For multi-speaker speech, we obtained one large significant positive cluster not only in left occipital cortex but also extending to left temporal regions (t_sum_ = 424,75, p < .001, d = .79). Descriptively, stronger tracking was found in right temporal regions, but this effect was not significant at a cluster-corrected level (t_sum_ = 45.25, p = .098, d = .48). We can show here that there seems to be some lateralization in the extent how visual speech is integrated and used for tracking the acoustic spectrogram. Already, despite not using any specific visual feature, increased tracking in occipital areas might indicate that correlated visual information is used in speech processing. We can observe this visual benefit strongly in multi-speaker speech where it extends to temporal regions which indicates processing beyond the visual system.

#### Added Value: Lip Movements

To investigate how the addition of a visual feature to the model increases prediction accuracy beyond acoustic tracking, we compared the combined model Spectrogram + Lip with the baseline Spectrogram model (middle row, Figure 2). This revealed significant increases in clear speech (t_sum_ = 156,50,p = .001, d = .65) and multi-speaker speech (t_sum_ = 216.06, p < .001, d = .68) in occipital areas compared to the spectrogram-only model. There was no additional visual benefit beyond the acoustic model in clear speech, but in multi-speaker speech (t_sum_ = 214.67, p < .001, d = .61). This highlights again the importance of visual information when acoustic information is unclear. While in clear speech, the increases in tracking over occipital areas might be simple responses of the visual system to visual information, a visual benefit (i.e. AV - A condition) is only present in multi-speaker speech.

#### Added Value: Lexical Units

Apart from our past research, other studies also investigated the benefit of lexical units in AV speech tracking (O’Sullivan et al., 2021). However, here we only found a significant increase in clear speech over right (t_sum_ = 306.69, p < .001, d = .50) and left temporal regions (t_sum_ = 295.29, p <.001, d = .61) as depicted in the bottom row of Figure 2. In contrast to the mentioned previous studies (Haider et al., 2023; O’Sullivan et al., 2021), no visual benefits beyond acoustic tracking were found. In multi-speaker speech, we only observed positive trends in the right (t_sum_ = 49.61, p = .054, d = .48) and left (t_sum_ = 37.15, p = .095, d = .35) temporal regions. This might indicate that higher-level lexical units are used to a lesser extent in difficult listening situations than in clear speech. This is in contrast to the effects of lip movements which occur especially in multi-speaker speech.

### Decreasing hearing ability leads to worsening of speech tracking in masked and audio-only conditions

As we now have a sample of an aging population with varying degrees of hearing ability, we wanted to investigate how individuals differ in their speech tracking as a measure. We especially focussed on differences in how individuals profit from visual speech and how they are affected by face masks. All other results are shown in Supplementary Material S2 and Supplementary Material S3. We can show that in general, speech envelope tracking of individuals with worse hearing ability is affected more when the speaker is wearing a face mask (*b* = 0.004, 95% CI = [0.001, 0.007], Figure 3A). To further illustrate this result, we split up the data into AV and A conditions and ran two separate models. We found a threefold interaction for the AV model between *Mask*, *Multispeaker,* and *HearingAbility*, which was absent in the A model (AV: *b* = 0.004, 95% CI = [0.000, 0.008]; A: *(b* = 0.000, 95% CI = [-0.004, 0.005], see Figure 3A).

**Figure 3.**
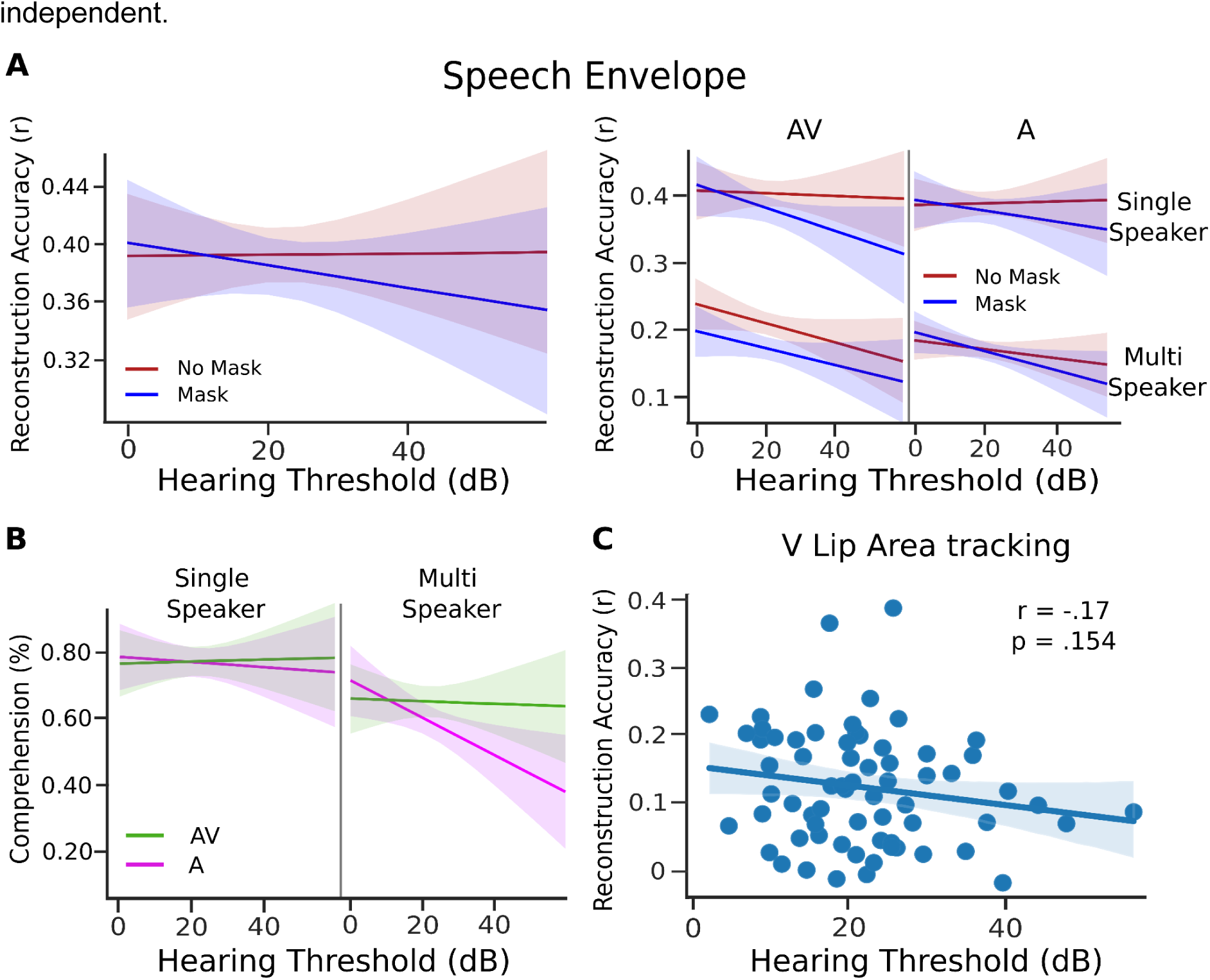
Effects of hearing ability on neural speech tracking and behavioral performance. **A** shows the reconstruction accuracy of the speech envelope as a function of hearing ability and if the speaker is wearing a face mask. On the left, the overall effect is shown, on the right the effect is split for single and multi-speaker speech. **B** depicts how hearing ability influences behavioral performance split up for single and multi-speaker speech in A and AV conditions. Here, the effect of hearing ability is driven by multi-speaker conditions in A speech. Please note the limitations of this effect described in the text. **C** describes the relationship between hearing ability and reconstruction accuracy of the lip area in V speech.

This might reveal that individuals with impaired hearing are using visual speech already in clear speech situations. However, as this effect is absent for the AV vs A contrast, acoustic degradation through wearing a face mask might also play a role here (Supplementary Material S2).

In their behavioral performance, participants with decreased levels of hearing ability are worse only in A conditions when there is a second speaker present (*(b* = 0.002, 95% CI = [0.000, 0.004], Figure 3B). Surprisingly, the face mask did not have an effect here (Supplementary Material S3). However, these results must be taken carefully, as in most conditions we only probe participants in two trials. As the response format has a 50% guess chance these results might prove unreliable. The presented evidence has proven inconclusive here. We can not directly say if the worsening of tracking or behavioral performance is a direct effect of visual speech or in the case of the face mask an effect of acoustic degradation. To investigate if individuals with decreased hearing are more proficient in extracting visual information, we calculated the reconstruction accuracy of lip movements in V conditions. Surprisingly, there was no correlation of hearing ability with the visual tracking score (r(67) = −.17, p = .154, Figure 3C). We therefore concluded that the ability to profit from visual speech does not have a relationship with individual hearing ability, but is independent.

### Tracking in visual-only speech is predictive for improved tracking in multi-speaker speech

Following this path, we analyzed if this visual speech tracking score (which is computed based on the V condition) is predictive of how individuals benefit from lip movements in AV speech. We first hypothesized that visual tracking in general is larger in multi-speaker than in single-speaker speech as shown in multiple previous works (Crosse, Di Liberto, & Lalor, 2016; Golumbic et al., 2013; Haider et al., 2023). We therefore tested this with a one-sided t-test (Figure 4A, left panel). The results show an increase in visual lip area tracking in the presence of a second speaker (t(66) = 2.08, p = .021, d = .51). In addition to confirming the previous results, Reisinger et al. (2023) and Aller et al. (2022) showed strong inter-individual differences in how visual information is used. We followed this up with an analysis of our visual tracking score as a between-subject level predictor (Figure 4A, right panel). Unsurprisingly, the visual tracking score (i.e. tracking of the lip area in V speech) is strongly predictive of how the lip area is tracked in AV speech *(b* = 0.502, 95% CI = [0.391, 0.610]). More interestingly though, this visual tracking score shows an interaction with the factor *Multispeaker (b* = −0.120, 95% CI = [−0.196, −0.044]). This shows that there are inter-individual differences in how people tend to or are able to use visual speech, and this comes especially into play in multi-speaker situations.

**Figure 4.**
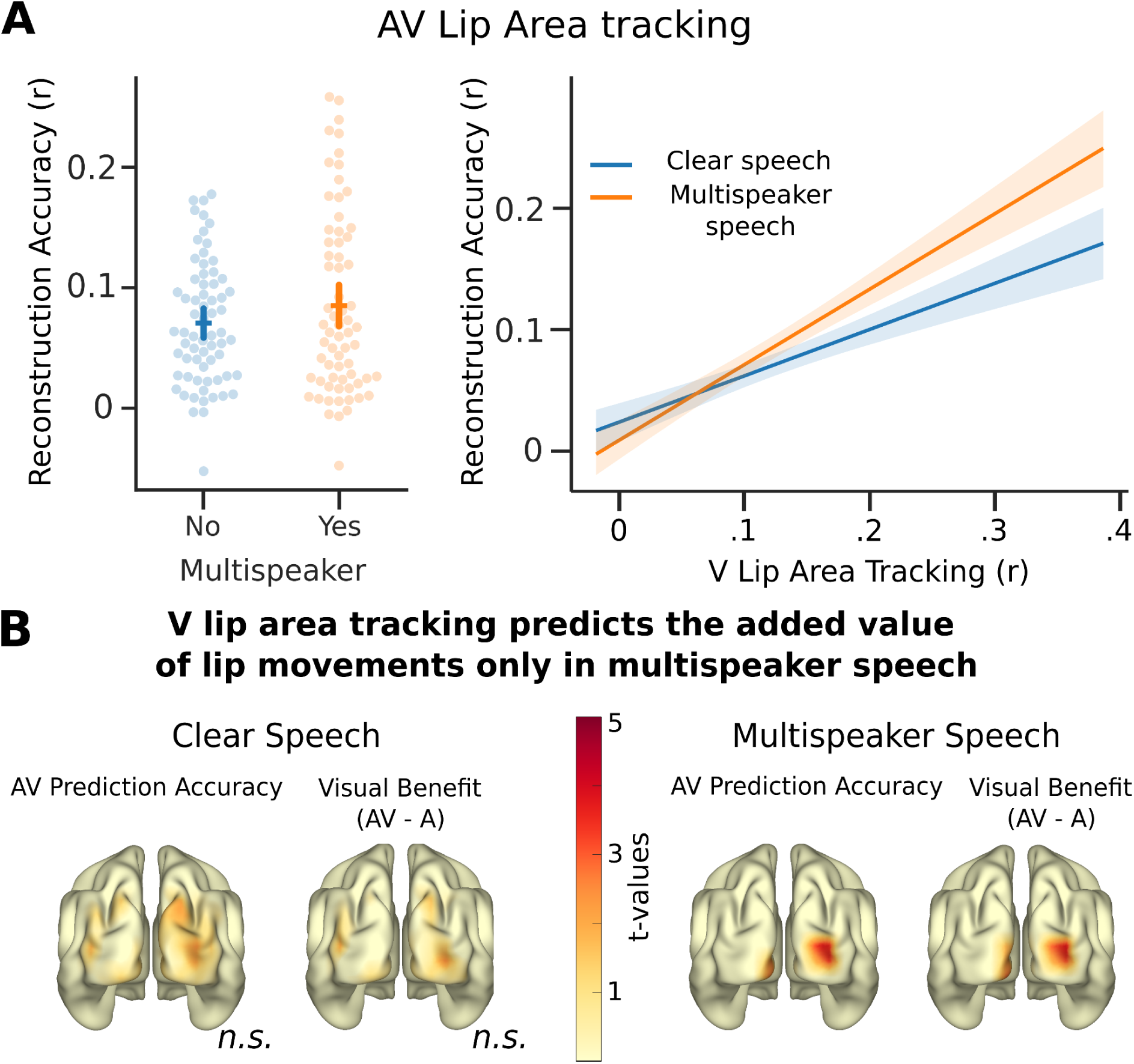
Visual speech benefits are stronger in AV multi-speaker speech and predicted by V tracking. **A** shows visual benefit based on backward modeling of lip movements. In the left panel, reconstruction of the lip area improves in multi-speaker compared to single-speaker speech. On the right, we observe how much individuals profit from visual speech - especially in multi-speaker situations - is predicted by their ability to track lip movements in V speech (silent videos). **B** highlights that V lip area tracking is correlated with the benefits of lip movements in occipital areas in multi-speaker speech, but not in single-speaker speech. For multi-speaker speech only significant and trend level voxels are shown.

We followed up this analysis in the forward model direction to localize these differences. We therefore computed the difference between the Spectrogram + Lip model and the baseline Spectrogram model (see Figure 2 *Added Value: Lip*). We restricted our analysis to the occipital cortex (168 voxels), as these were the only regions showing added value from lip movements (Figure 2). We then used the Spearman correlation between each voxel and the individual visual speech tracking score. Afterward, we used the same cluster-correction method as before with 10000 permutations. The results of this analysis are shown in Figure 4B.

By using this method, we found no cluster in clear speech neither for the contrast between the AV conditions nor for visual benefit. This is in contrast to the effects found in multi-speaker speech. Here we found one significant cluster for the AV prediction accuracies in the right occipital cortex (t_sum_ = 60.06, p = .025, r = .30) and a trend in the left occipital cortex (t_sum_ = 39.75, p = .056, r = .26), as well as single cluster for visual benefit extending to both hemispheres (t_sum_ = 102.17, p = .003, r = .28). These results show that people indeed differ in their ability or tendency to extract visual information. As proposed by inverse effectiveness individuals seem to mostly recruit this information when acoustic information is noisy.

## Discussion

In this manuscript, we wanted to extend previous findings in the speech tracking domain to an aging population with varying degrees of hearing ability. We can mostly replicate previously found effects in young (effectively student) populations. Especially, we can again show the importance of visual speech in multi-speaker speech, meaning in situations where acoustics are unclear. However, results regarding the impact of hearing ability are mixed. Here, we expected a stronger reliance on visual speech in individuals with decreasing hearing ability. Despite finding adverse effects in envelope tracking through a face mask in these individuals, we cannot pinpoint these results to missing visual information. Although we observe this effect of the face mask to be mostly driven by AV modalities, the threefold interaction of the interaction of *Mask*, *Modality,* and *HearingAbility* proved to be not significant, therefore we can not attribute this effect to missing visual speech directly. Individuals with decreased hearing ability may be impacted by face masks in two ways. First, through degraded acoustics, as face masks reduce clarity in higher frequencies (Corey et al., 2020), which does not have an impact on normal-hearing individuals. And second by missing visual information which affects both populations. We expected hearing-impaired individuals to be more proficient and reliant on this visual information. However, our results show no correlation between hearing ability and visual tracking score (i.e. lip area tracking in V speech), and thereby the notion that individuals get better at using visual speech as they more strongly rely on it, was not supported. The visual tracking score predicts how individuals use visual information in AV speech, especially in multi-speaker situations when acoustics are unclear.

### The added value of lip movements is especially important in multi-speaker situations

In previous studies, it has been shown that speech tracking is enhanced in AV speech compared to A-only speech (Crosse, Di Liberto, & Lalor, 2016; Haider et al., 2023; O’Sullivan et al., 2021). Using an almost identical paradigm to the present study, we confirmed this benefit of visual speech in two previous studies, however with some differences in the specific effects. In the study by Reisinger et al. (2023), a strong benefit of lip movements in multi-speaker speech was found, while it was absent in Haider et al. (2023). While the current study is an experimental continuation of Haider et al. (2023), our results point more in the direction of Reisinger et al. (2023) and other previous investigations (Crosse, Di Liberto, & Lalor, 2016; Crosse et al., 2015). Also, our current results are in line with classical behavioral findings about visual speech, which highlight their importance in multi-speaker situations (Sumby & Pollack, 1954). These differences between our studies might be explained by a caveat of speech tracking paradigms in general: As all speech features (visual and acoustical) are strongly correlated (Chandrasekaran et al., 2009), it is not easily possible to disentangle visual and acoustic tracking completely. This means, that when we investigate tracking of the speech spectrogram correlated visual features (i.e. lip movements) alongside acoustic information can be deduced from the signal. Thereby, tracking of visual information in occipital areas is already observed at this stage. By adding lip movements to the model, we can further increase the amount of visual information modeled by the TRFs. However, how much visual information was tracked before is unclear and might vary over subjects and conditions. Therefore, the increase of information through lip movements is varying.

Despite this, the picture that visual information is beneficial for speech tracking, especially in situations with unclear acoustic (e.g. by adding a second speaker or white noise) is confirmed in many studies. What is debatable, is if and how tracking lexical units like phonemes benefit from visual speech. Studies by O’Sullivan and colleagues (2021) and Nidiffer and colleagues (2023) show some evidence that phonemes are tracked better in the presence of visual speech and are tracked directly in visual areas. However, our two studies Haider et al. (2023) and the current one do not confirm this. One must note here the differences used in the experimental design and analysis which might give rise to these inconclusive findings. Taken together, we can demonstrate visual benefits in speech processing through lip movements. The working mechanism here might be on the one hand the access to temporal cues as proposed by Peelle and Sommers (2015). On the other hand, we cannot provide neural evidence for the second mechanism proposed by the model, which is the direct extraction of visual information for phoneme discrimination (i.e. the added visual benefit of lexical units).

### Neural speech tracking in individuals with decreasing hearing ability is affected by face masking

There have been several studies in the past investigating speech tracking in the hearing-impaired population. One such study by Gillis et al. (2022) found an increased and delayed response to speech in individuals with hearing loss compared to normal-hearing individuals. Another study found that increased tracking was associated with better speech comprehension in hearing-impaired individuals (Schmitt et al., 2022). While several studies investigated speech tracking in the context of visual benefits, only one study to our knowledge investigated potential visual benefits in individuals with decreased hearing ability (Puschmann et al., 2019). In general speech tracking scores seem to be not affected by hearing ability, but individuals with worse hearing are affected more by a face mask mainly in AV conditions. As others have pointed out in behavioral studies, these individuals profit the most from visual speech and therefore it is understandable that they show decreases in speech tracking through masking (Brown et al., 2021). However, this effect cannot only be attributed to missing visual speech in these individuals. While our previous study with young normal-hearing individuals could show that a face mask only has an effect in AV conditions, here the findings are more mixed. There is some indication that the effect is stronger AV compared to A conditions (Figure 3), but not significantly. Previous studies showed acoustic degradations of speech features for speakers wearing a face mask (Corey et al., 2020; Rahne et al., 2021). Despite these mostly being restricted to higher frequencies, this might have an impact on listeners who have trouble hearing and need every information available to aid their speech processing (Motlagh Zadeh et al., 2019). Therefore, the effect of a face mask on the hearing impaired might point to a combined effect here. It is important to view these results in light of our experimental setup. We did set the volume of the stimuli presentation to a comfortable level at the start of the experiment. This might lead to an underestimation of the effect hearing ability has on speech tracking.

Contrary to previous assumptions, we did not find stronger visual tracking with declining individual hearing. Intuitively, as hearing decreases a stronger reliance on visual speech in everyday conversations leads to stronger proficiency or a tendency to track this information. This seems not to be the case. We find that there is no correlation between hearing ability and visual speech tracking (i.e. tracking of lip movements in silent videos) in general. Therefore, we propose that the ability to extract visual information from lip movements is a trait independent of an individual’s hearing ability. Affected individuals might use a wide range of compensatory mechanisms not only involving visual speech, from hearing aids to cognitive compensation (Başkent et al., 2016).

### Visual speech tracking is predictive for visual speech benefits primarily in multi-speaker speech

We can confirm our previous results showing increased visual benefits in multi-speaker speech (Reisinger et al., 2023). Additionally, as that study and Aller et al. (2022) noted before, there seems to be a strong interindividual difference in how people use visual speech. Some people do not seem to use visual information at all, not even in multi-speaker speech, while others profit greatly from it. We here firstly demonstrate that this interindividual difference can be measured in V speech and that this score is predictive of benefits in a more naturalistic AV speech. It has to be noted that this finding must be confirmed by behavioral results in other studies, as the quality of our comprehension measures is quite low (as described in the results section). However, if it holds true, this offers theoretical and other methodological implications. Regarding theoretical implications, we must differentiate between individuals profiting from visual speech and others who do not. This might include adaptive measures in individuals who are hard of hearing to the individuals’ specific needs. Some individuals might use more visual information, some might use cognitive compensation (Başkent et al., 2016). The latter behavior however is associated with increased listening difficulty and therefore also associated with social withdrawal (Hughes et al., 2018). If in general, it is possible to train the ability to track visual speech, this might be a public health measure in an aging population. Again these findings must be confirmed by strong behavioral studies taking into account comprehension performance and listening difficulty.

### Future directions

With this study, we open interesting new opportunities to analyze neural data in the context of AV speech. We can present a measure that might serve as a trait to determine individuals’ visual speech tracking abilities. However, it must be evaluated how meaningful this measure is behaviourally. There is some evidence by (Reisinger et al., 2023), but further investigation is needed. Also speech tracking in the context of hearing loss, must be evaluated using a combination of strong behavioral measures and neural measures. So far, it is not easily possible to link speech tracking to comprehension performance. Finding this link between neural scores and behavior opens exciting new possibilities to evaluate different acoustic and visual features in normal-hearing and hearing-impaired individuals. Lastly, a systematic investigation of different experimental designs must be conducted. So far we cannot rule out that unsystematic differences between experimental setups are responsible for the found differences between labs (e.g. O’Sullivan et al., 2021), but also within a research group (e.g. Haider et al., 2022, 2023).

## Conclusion

We can confirm the previous results on neural speech tracking and the added value of specific visual features (i.e. lip movements). We extend a previous study by our lab in sensor space and confirm the proposed underlying brain regions. We also show that individuals with hearing loss are indeed more affected by face masks concerning speech envelope tracking, but still, it is unclear in this population if this effect is produced only by blocking visual speech or by both visual speech and acoustic degradation. This might have important implications for using face masks in critical environments like hospitals (Chodosh et al., 2020). Lastly, we show that people with decreasing hearing do not necessarily become better at extracting compensating visual information. This ability seems to be an individual trait and importantly predicts how an individual uses visual speech in the context of degraded acoustics.

## Code Availability Statement

The code to reproduce statistical analyses can be found at https://gitlab.com/CLH96/visualspeechtrackinghearing. Further code can be supplied upon request.

## Author Contribution (CRediT)

**Chandra Leon Haider:** Conceptualization, Methodology, Software, Validation, Formal analysis, Investigation, Data Curation, Writing - Original Draft, Writing - Review & Editing, Visualization, Project administration.

**Anne Hauswald**: Conceptualization, Writing - Review & Editing, Supervision.

**Nathan Weisz**: Conceptualization, Resources, Writing - Review & Editing, Supervision, Funding acquisition.

## Acknowledgments

Sound icon made by *Smashicon* from www.flaticon.com.

I would like to thank the whole research team and especially Fabian Schmidt for always giving good advice about methods, figure design, and all in all great insights.

## Supplementary Material

**Supplementary Material S1.**
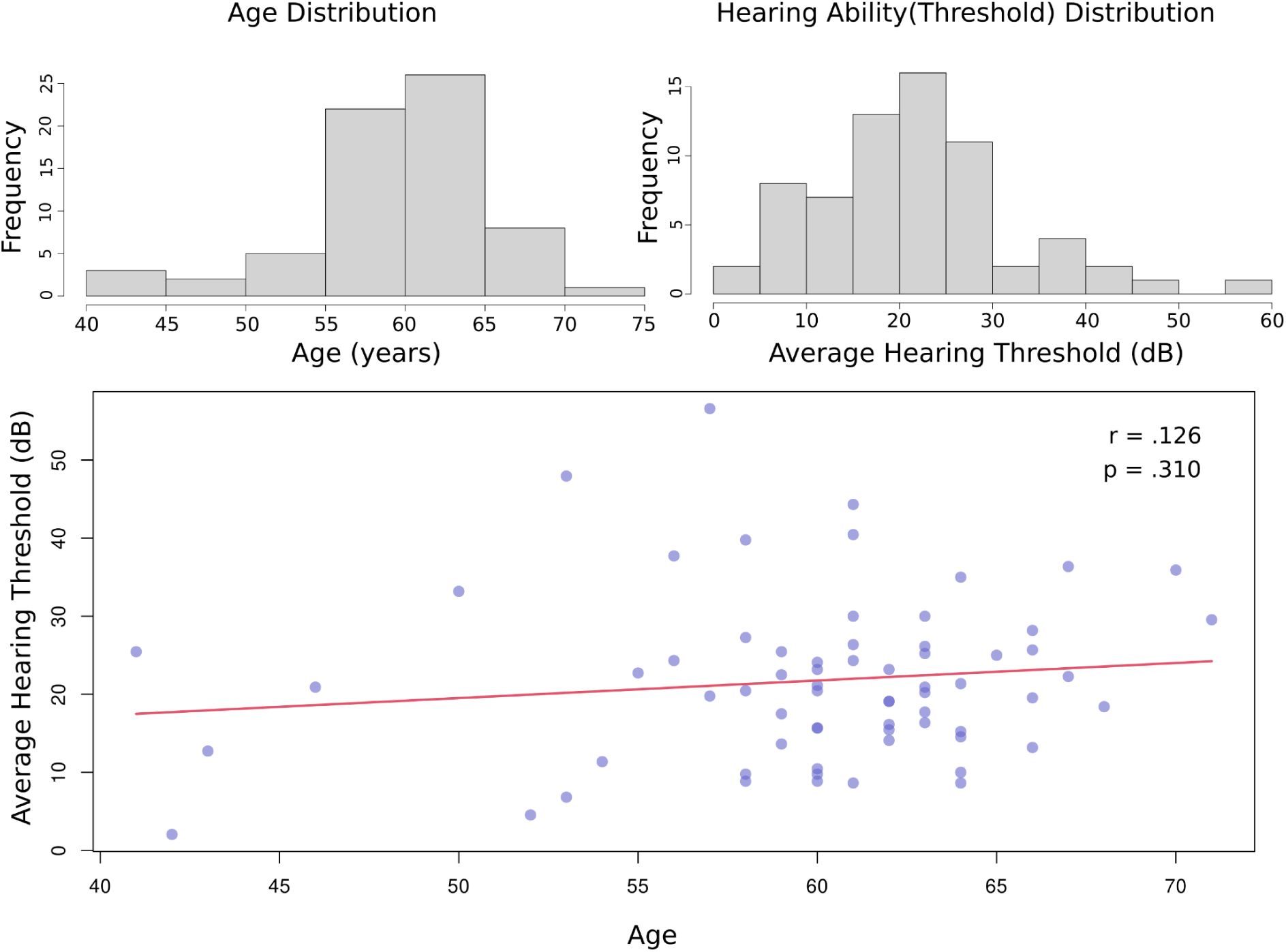
Age and Hearing ability information. This figure shows the age distribution as well as the hearing ability (i.e. individual hearing thresholds averaged across ears and frequencies). We furthermore show that in our sample hearing ability and age are uncorrelated, which allows investigation of hearing ability irrespective of age.

**Supplementary Material S2.**
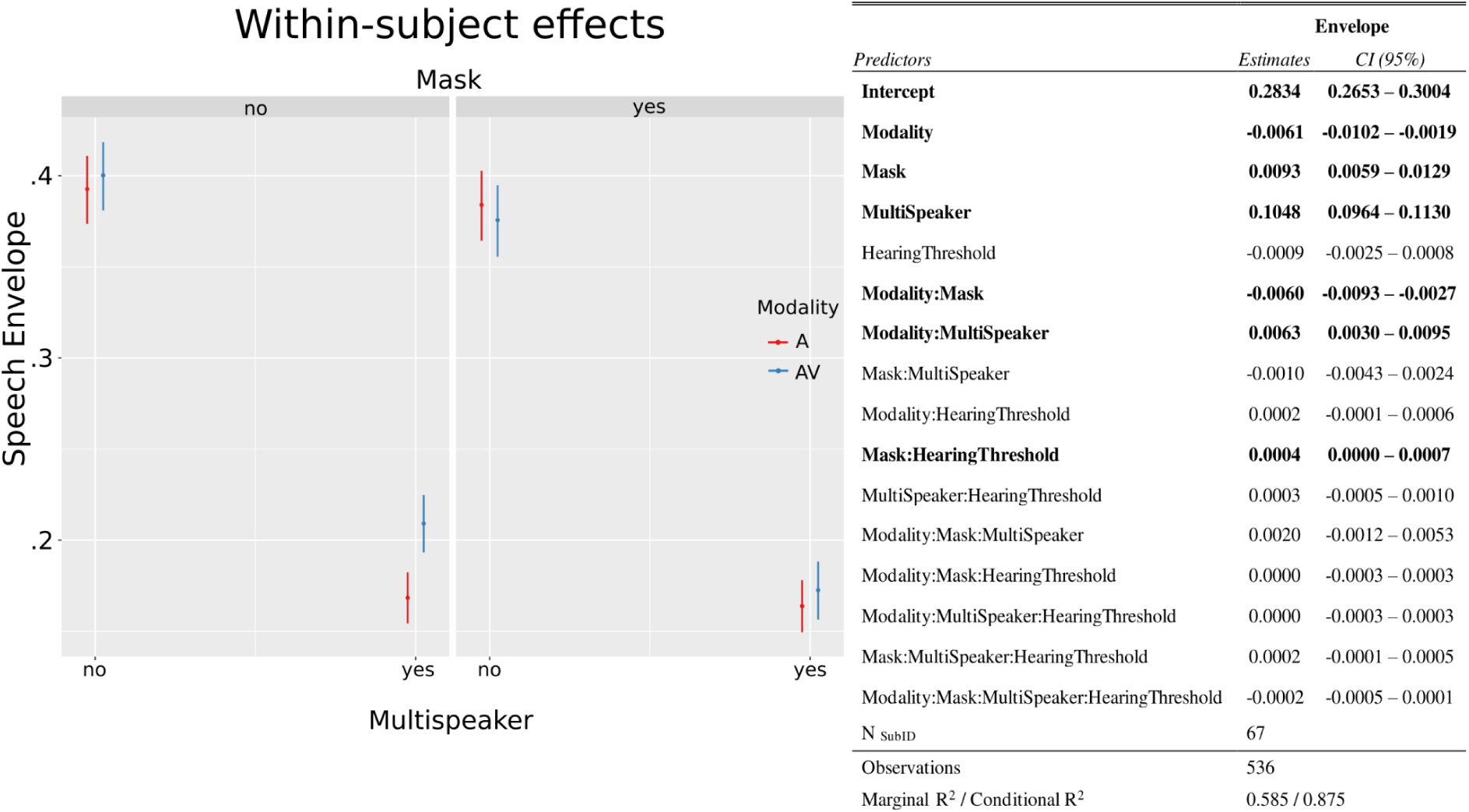
Results of the reconstruction of the speech envelope. On the left, we show the within-subject effects (i.e., our experimental manipulation) of the reconstruction of the speech envelope. For that purpose, the between-subject factor (i.e., HearingAbility) was centered. The corresponding statistics can be seen on the right-hand side. Significant effects are shown in bold.

**Supplementary Material S3.**
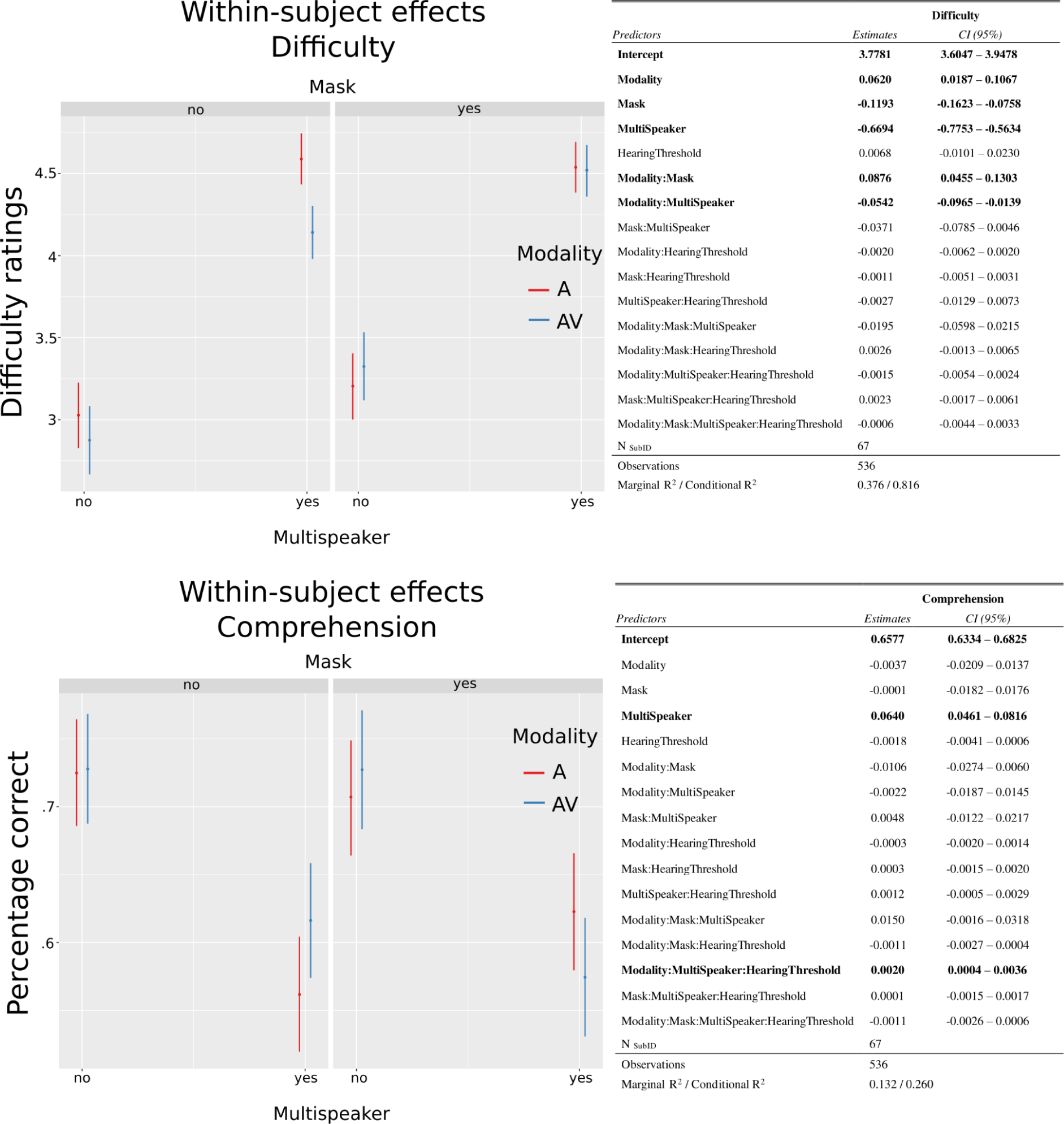
Results for Comprehension performance and difficulty ratings. On the left, we show the within-subject effects (i.e. our experimental manipulation) for difficulty ratings and comprehension performance. For that purpose, the between-subject factor (i.e. HearingAbility) was centered. The corresponding statistics can be seen on the right-hand side. Significant effects are shown in bold.

